# Simulations show increased lipid interdigitation in transmembrane coupling of protein condensates

**DOI:** 10.64898/2026.07.06.735898

**Authors:** Ketsia Zinga, Jeanne Stachowiak, Pengyu Ren

## Abstract

Liquid-liquid phase separation of proteins has been observed to occur on biological membranes, where it is thought to play a role in diverse cellular behaviors. Recent work has demonstrated colocalization between protein condensates on opposing leaflets of the bilayer, suggesting that protein phase separation may be coupled across the bilayer. However, the mechanism behind this coupling phenomenon remains poorly understood. Here we seek to understand the protein-protein and protein-membrane interactions that give rise to transbilayer coupling of protein condensates. We perform coarse-grained molecular dynamics simulations of a bilayer with a disordered protein condensate tethered to each leaflet surface. In this system, we observe stable, coupled diffusion of the condensates across the membrane. We find that increasing the protein-protein interaction strength leads to decoupling, driven by competing membrane curvatures induced by each condensate. However, by applying membrane tension we suppress curvature and restore coupling even at higher protein interaction strengths. Under coupling conditions, we find that lipid entropy is reduced upon direct contact with proteins, but this effect is not transferred to the opposing leaflet. Interestingly, further analysis reveals increased transverse lipid packing (interdigitation) beneath the condensates relative to protein-free regions. Based on these observations, we propose that enhanced lipid interdigitation mediates interleaflet communication and serves as the primary mechanism driving transbilayer coupling of condensates in this system. This work provides insight into a potential physical mechanism for transmembrane communication in cellular contexts and suggests directions for future investigation.

**Significance Statement:** Liquid-like condensates are active participants at cellular membranes, where they act as organizers and catalysts for various cellular processes. Recent work has demonstrated that protein condensates can couple across the bilayer; however, the molecular mechanism of this transbilayer coupling remained unknown. Here, we investigate the molecular basis of transmembrane condensate coupling through detailed analysis and propose a mechanism for the phenomenon. This work advances our understanding of how information is transmitted across the bilayer, with implications in cellular requiring coordination across the membrane, such as signaling, and more broadly in the field of membrane biophysics.

## Introduction

Liquid-liquid phase separation (LLPS) has emerged as an important organizing principle in cells, in contrast to the traditional paradigm of membrane-bound compartmentalization^1–4^. LLPS of biomolecules occurs in both the cellular cytoplasm and nucleus. LLPS arises from weak, transient, and multivalent interactions among proteins, which often contain intrinsically disordered regions (IDRs) ^5,6^. The resulting liquid-like assemblies, or condensates, are characterized by rapid molecular exchange with the environment and high surface tension which allows them to merge and re-round^3^. Through these properties, condensates participate in diverse cellular functions, including regulation of RNA transcription, stress responses, and signal transduction^7–13^.

It is increasingly evident that condensates are active at membrane surfaces. These assemblies have been observed at the plasma membrane, organelle membranes, and intracellular membrane contact sites, where they contribute to the spatial organization of signaling and trafficking machinery^8–10,14–16^. For example, the early endocytic adaptor proteins Fcho and Eps15, which are involved in clathrin-mediated endocytosis (CME), form condensates whose liquid-like properties are important for endocytic pit progression^14^. Similarly, liquid-like condensates of actin associated proteins such as N-WASP and VASP can nucleate and polymerize actin filaments, a process essential for propagation at the leading edge of migrating cells and cellular signaling^15–19^.

The association of protein condensates with membrane surfaces is not unexpected, as confinement to a two-dimensional surface concentrates proteins and lowers the critical concentration required for phase separation compared to the three-dimensional bulk^8,14,20^. Additional studies demonstrate that condensate interactions with lipid bilayers can influence membrane organization and signaling. One such example is the T-cell signaling complex. Condensates of the transmembrane protein linker for activation in T-cells (LAT) and two additional signaling molecules, Sos and Grb, are capable of stabilizing and inducing the formation of lipid domains in reconstituted systems^21^. In turn, ordered lipid domains both localize and stabilize LAT/Sos/Grb condensates in both reconstituted and cellular systems. In addition to T-cell signal transduction, several other membrane-associated cellular processes which feature LLPS coordinate protein assemblies on opposing leaflets of the bilayer, such as membrane trafficking events and cell-cell communication^9,22–25^.

Recent studies have demonstrated that condensates of intrinsically disordered proteins can couple across lipid bilayers^26,27^. When phase-separating proteins are present on both sides of the membrane, condensates form independently on each leaflet but frequently become spatially aligned across the bilayer. Once established, these paired condensates remain colocalized over several minutes. Despite the robustness of this phenomenon, the molecular basis behind this transbilayer coupling remains poorly understood.

In this work, we investigate the mechanisms by which intrinsically disordered protein condensates couple across lipid bilayers. Using coarse-grained molecular dynamics simulations with the Martini force field, we model condensates of RGG domains tethered to both sides of a lipid bilayer.

In our simulated system, we observe that condensates of RGG recapitulate the stable coupling behavior observed *in vitro*. We quantify persistent overlapping area between the two condensates and correlated diffusion under several conditions. We then perform a detailed investigation of the lipids in the system in order to identify a mechanism of information transfer through the bilayer. This work advances understanding of transbilayer communication and provides insight into how protein assemblies interact through cellular membranes to coordinate membrane-associated biological processes.

## Results and Discussion

### Condensates in the simulation demonstrate coupled diffusion through the bilayer

In order to investigate the mechanism behind transbilayer coupling of condensates, we modeled a protein-bilayer system utilizing the Martini 3.0 coarse grain force field^28,29^. We chose to model condensates of the RGG-rich domain of LAF-1 (RGG), a well-characterized protein model for the study of LLPS on membranes and that was employed in previous studies of transbilayer coupling of condensates^11,26^. We began by placing two pre-formed condensates each containing 32 individual RGG chains on opposing leaflets of a 50 x 50 nm^2^ POPC bilayer. The condensates were positioned such that their projections in the plane of the membrane (xy plane) overlapped partially (Figure 1A and Supplemental Figure 1A, first panel). Previous experimental studies of protein condensates on lipid bilayers utilize Ni-NTA his-tag chemistry to attach proteins to the bilayer surface. To emulate this bond in our system, we tethered each protein to the bilayer using a constant force between the center of mass of the first two N-terminal residues and the nearest lipid head group at initialization. This force was maintained for the duration of the trajectory. We then solvated the system at 100 mM NaCl using the *insane* program^30^. The Martini force field overestimates protein-protein interactions, especially for disordered proteins^31–34^. In a previous work, we identified a scaling factor of α = 0.98-0.975 applied to protein-protein (van der Waals) interactions in order to capture the liquid-like nature of RGG condensates with the Martini force field^35^. Therefore, we began our study with α = 0.98, which presented both a well-defined condensed and a dilute phase for RGG. Following minimization and equilibration, we performed a 10 µs production simulation and two additional 15 µs repeat production simulations. Details for membrane and condensate generation, minimization, equilibration, and production simulations are further described in methods.

**Figure 1.**
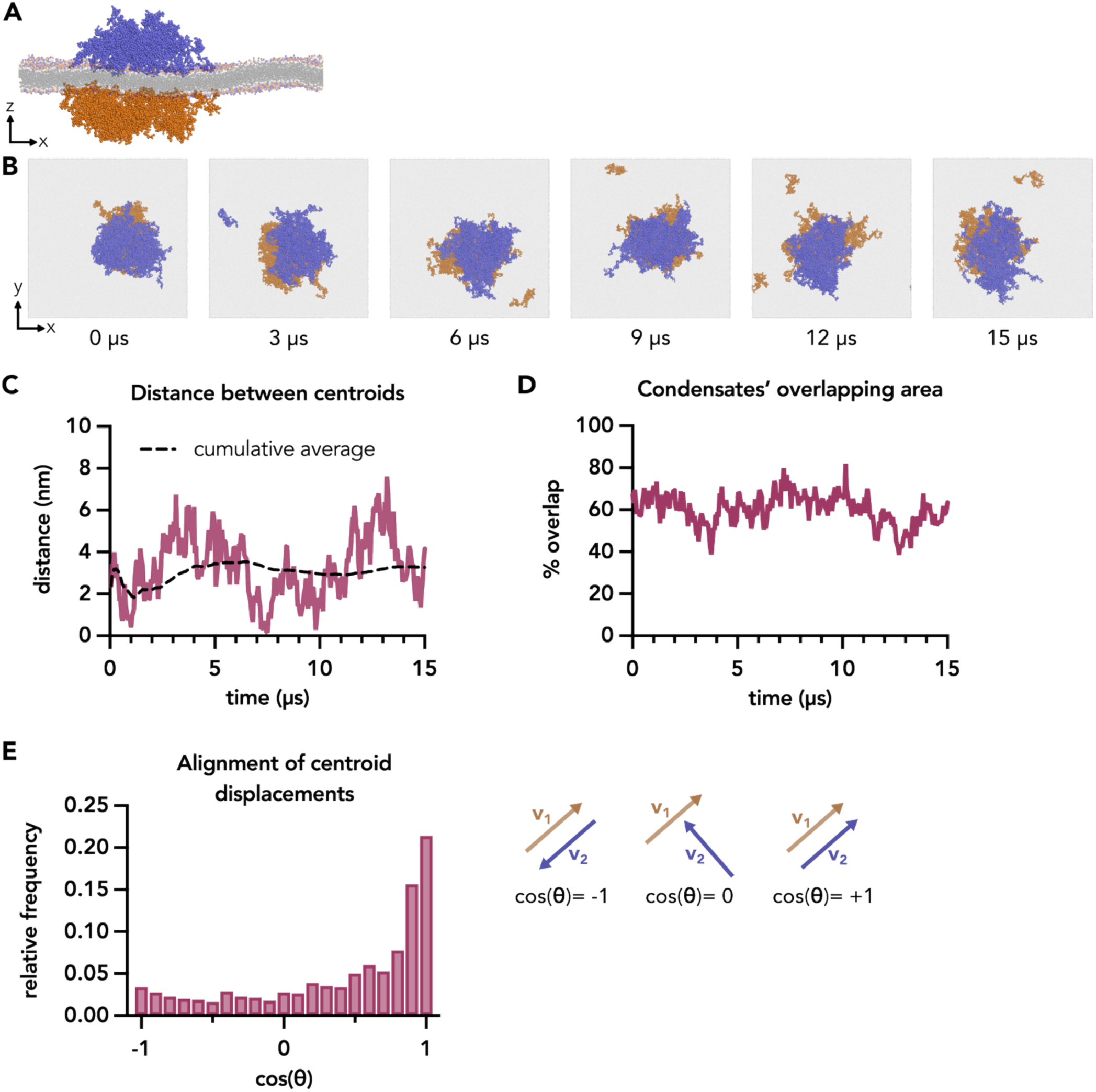
RGG condensates in the simulation demonstrate coupled diffusion across the bilayer. **A.** Cross-sectional view of the condensates and membrane. **B.** Representative snapshots shown from above the xy plane with the bilayer rendered in translucent grey, upper leaflet proteins colored blue, and lower leaflet proteins colored orange. A condensate was placed on the surface of each bilayer leaflet such that their projections were overlapped. The condensates diffuse together throughout the trajectory while 1-2 protein chains sample the dilute phase. **C.** 2D distance in the plane of the bilayer (xy plane) between condensate centroids and the cumulative average over the trajectory shown in A. The distance varies from 0 to over 6 nm, while the average remains near 3.3 nm throughout the trajectory. **D.** Overlap of the xy-projected area of the condensates shown in A, expressed as a percentage of the total condensate area over time (see equation 1). **E.** Distribution of the alignment of in-plane (xy) displacements of the upper and lower leaflet condensate centers of mass, with accompanying illustrations. Values range from −1 (antiparallel) to 0 (orthogonal) to +1 (parallel, aligned). Data was grouped into 21 bins with centers at increments of 0.1 units. Data displayed was taken from three independent trajectories totaling 40 µs.

Representative snapshots from a trajectory are shown in figure 1B, viewed from above the xy plane. Proteins tethered to the surface of the lipid bilayer’s upper leaflet were colored blue, proteins tethered to the surface of the lipid bilayer’s lower leaflet were colored orange, and the bilayer was colored gray and rendered translucent for clarity. Over the course of the trajectory, we observed that the condensate on the top leaflet (blue) and the condensate on the bottom leaflet (orange) maintained a large degree of overlap for the duration of the 15 µs trajectory and appeared to diffuse synchronously. Up to two individual protein chains were observed to dissociate from the main condensate at a time (Figure 1B, panels 2-5). This sampling of the surrounding space was indicative of LLPS, in which proteins dynamically exchange between an enriched, condensed phase or a depleted, dilute phase.

To characterize the degree of transbilayer coupling in the system, we first quantified the distance between the two condensates’ centroids (Figure 1C). Despite demonstrating coupled diffusion and generally high overlap percentage, the distance between the condensates was generally non-zero throughout the trajectory. Though some noise was expected due to shifts in the amorphous condensate boundaries, the average distance converged to ∼3 nm. This data suggested that the condensates were stably coupled at a slightly offset conformation, rather than exactly aligned in space.

To further characterize the condensates’ relative positioning, we next quantified the percent overlap as below:

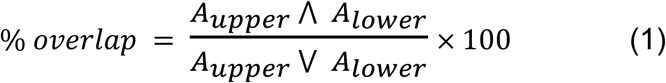

Where A_upper_ and A_lower_ are the areas covered by the upper and lower leaflet condensates, respectively, calculated by projecting the protein particles’ positions onto a grid in the xy plane. Protein chains which were not directly in contact with the primary protein mass were considered as the dilute phase and not included in the calculation. We took the sum of grid spaces occupied by *both* the upper leaflet and the lower leaflet condensate (A_upper_ ⋀ A_lower_) to find the overlapping area. We then took the sum of grid spaces occupied by *either* the upper leaflet or the lower leaflet condensate (A_upper_ ⋁ A_lower_) to find the total protein-occupied area. The overlapping percentage was then calculated by dividing as in Equation 1. Over the trajectory shown in figure 1B, the overlapping area oscillated around an average value of 60% (Figure 1D). The maximum observed overlap did not reach 100%, but is determined by both the offset of the condensate centroids and their irregular shape as seen in figure 1A.

These analyses were performed for two additional repeats (Supplementary figure 1A-1D). Overall, stable coupling was observed for two out of three repeats. However, in repeat 3, after 10 µs, the distance between the centroids increased to 13 nm and the percent overlap dropped to less than 20%, indicating a “decoupling” of the two condensates (Supplementary figure 1C-1D).

In order to further quantify the degree of coupling, we introduced an alignment metric between the movement of the two condensates (Figure 1E). For each condensate, we obtained a time series of its in-plane (xy) center of mass coordinate at intervals of 50,000 ps. Protein chains which were in the dilute phase were not included in the calculation. From this time series, we then calculated the displacement vector between time points to obtain a series of displacement vectors for each condensate. For each timepoint, we then took the cosine of the angle between the upper and lower condensate displacement vectors according to equation 2 below:

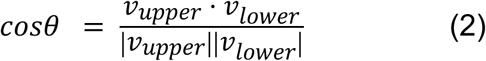

This value ranged from −1, when the displacements were completely anti-parallel, to 0 when they displace orthogonally, and 1, when the displacements were exactly parallel and considered fully aligned. We first validated the metric by calculating the alignment of a condensate to itself and of two condensates in separate, unassociated trajectories and found that the method yielded the expected distributions (Supplementary Figure 1E and 1F). We then calculated the alignment for all three trajectories and summarized as the normalized distribution in Figure 1E. Over 50% of displacements were well-aligned, with the angle (*θ*) between the two vectors less than 45 degrees (cosθ = 0.6). Approximately 40% of sampled timepoints fell within the upper two bins, corresponding to an angle of less than 30 degrees between upper and lower condensate displacement vectors. Overall, this data suggested that the coupled condensates diffused with an alignment greater than that which would be expected for two randomly diffusing condensates.

### Coupling of condensates is a local energetic minimum

To further investigate the apparent offset of the coupled condensates, we then explored the energetic landscape as a function of the distance between the condensate centroids (Figure 2). We used the Gromacs command *gmx distance* to obtain the xy in-plane distance between the upper and lower leaflet condensate centroids for all three repeat simulations performed at α = 0.98. We then calculated the probability distribution of centroid-to-centroid distances across all three repeat production simulations, as well as for each individual trajectory (Figure 2A). The second repeat exhibited the narrowest distribution, indicating sustained strong coupling throughout the trajectory (Figure 2B, yellow). Within the 10 µs window, its most populated bin was centered at a centroid-to-centroid distance of 6.75 nm. The first simulation showed a slightly broader distribution (Figure 2B, blue), although this trajectory began with partial overlap and may include an equilibration period prior to full coupling. In contrast, the third simulation, which shared the same initial configuration as repeat 2, displayed the broadest distribution (Figure 2B, green), with a maximum centroid-to-centroid distance of 18 nm, approximately twice that observed in the other two trajectories. By the end of this trajectory, the condensates appeared to decouple. That is, they no longer exhibited coupled diffusion or stable overlap.

**Figure 2.**
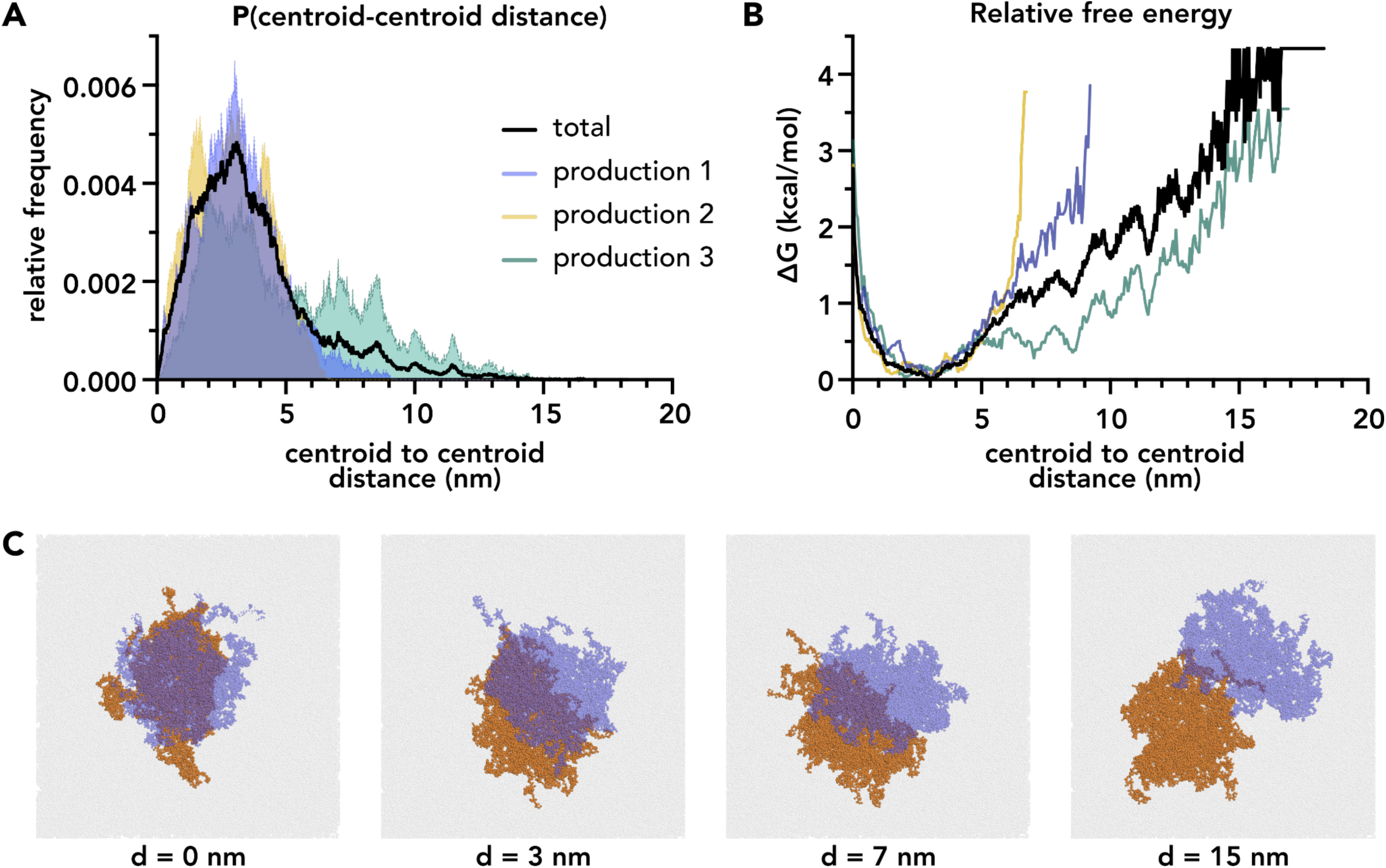
Coupling of condensates is a local energetic minimum. **A.** Probability distributions of distances between condensate centroids in three repeat production simulations, shown in black. Though production simulations two and three were performed for 15 µs, only distances from 0.05 to 10 µs of each simulation were used to maintain even weighting across repeats. Distributions for individual trajectories are also shown with production 1 colored blue, production 2 colored yellow, and production 3 colored green. Data was binned in intervals of 0.02 nm and normalized so that the sum of all bins equaled 1. **B.** Free energy surface of the xy distance between centroids in three repeat simulations (bold black) and for each of three repeat trajectories (coloring consistent with 2A) calculated using the probability distributions in 2A. **C.** Snapshots demonstrating condensate positioning across four centroid-to-centroid distances with the upper leaflet condensate (blue) rendered partially transparent to visualize overlap with the lower leaflet condensate (orange). The minimum distance was 0 nm (first panel). The energetic minimum was ∼3 nm (second panel). The relatively higher energy transition was around 7 nm (third panel). The distance at which condensates had little to no overlap was ∼15 nm (fourth panel).

From the probability distribution, we then calculated the relative Gibbs free energy, *ΔG*, for each distance as described in equation 3 below:

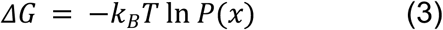

normalizing to the maximum frequency such that *ΔG* at *P*_max_ was zero.

The free energy surface calculated from all three trajectories revealed an energetic minimum at a centroid-to-centroid distance of 3.1 nm (Figure 2B, black), corresponding to ∼60-70% overlap. An energetic barrier of approximately 1 kcal/mol was observed for the separation to be greater than 6 nm. Each individual trajectory also exhibited almost identical energetic minimums at a distance of 3.1 nm. Further inspection of the two trajectories which remained well-coupled for their duration (Figure 2B, blue and yellow) revealed that the energy barrier can be as high as 4 kcal/mol between 6 and 10 nm separations.

Beyond 15 nm, the condensates exhibited little to no overlap (Figure 2C, fourth panel). Sampling of intermediate states between overlapping and complete separation was limited, particularly between 6 and 15 nm centroid-centroid distance. Despite the limited scope of our simulations, we concluded that within the explored energy landscape, the coupled state at ∼3 nm centroid-to-centroid distance functioned as a locally stable energetic minimum.

### The degree of condensate coupling is dependent on the strength of protein-protein interactions

The strength of protein-protein interactions within the condensate has been shown to influence its interaction with the bilayer. We therefore examined how the choice of α affected the observed coupling behavior. We chose to investigate α values of 0.9775 and 1 (unscaled) applied to protein-protein Van der Waal interactions. To initialize the new systems, we utilized an initial conformation with the highest degree of overlap observed from the simulations at α = 0.98. We performed a 20 µs simulation with α = 1.00 scaling, as well as three repeat simulations totaling 32 µs with α = 0.9775. Representative snapshots of the trajectories (Figure 3A and 2D) use coloring consistent with Figure 1.

**Figure 3.**
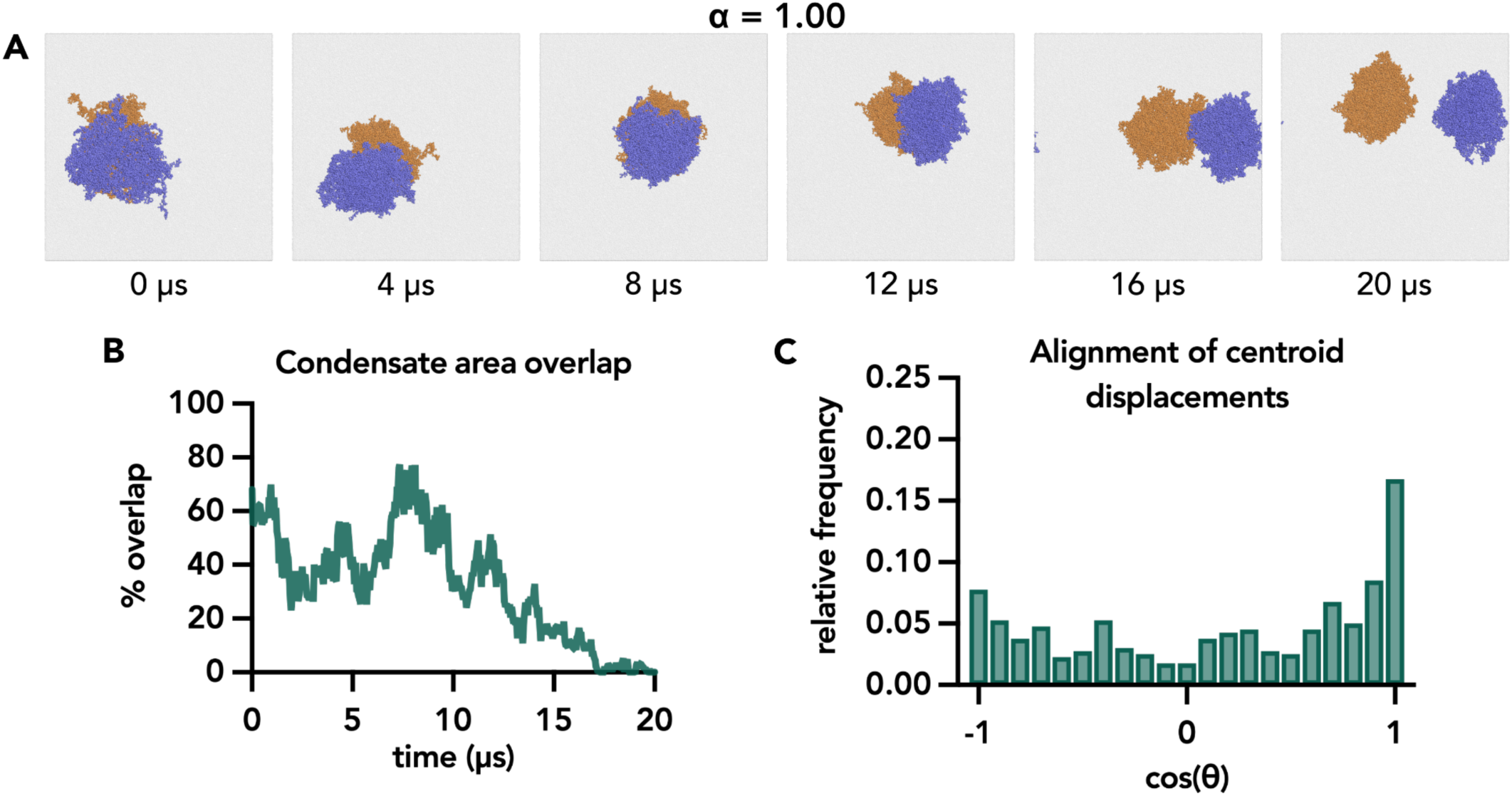
Suppression of membrane curvature and fluctuations rescues stable coupling when protein-protein interactions are increased. **A.** Time series snapshots shown from above the xy plane of a trajectory with increased protein-protein interactions with a scaling factor of α = 1.00. The condensates began with a high degree of overlap which was maintained for the first half of the trajectory. The condensates moved apart in the later portion of the trajectory until they were no longer overlapping or diffusing in a coupled manner, **B.** Overlap of the xy-projected area of the condensates in A expressed as a percentage of the total projected condensate area over the course of the trajectory in A. The overlapping area decreased after 10 µs until reaching 0 %. **C.** Distribution of the alignment of in-plane (xy) displacements of the upper and lower leaflet condensate centroids for the trajectory in A with increased protein-protein interaction strength. Data was grouped into 21 bins with centers at increments of 0.1 units.

When the α = 0.9775 scaling factor was applied, we observed expansion of the protein condensates and more amorphous geometry compared to α = 0.98, a result of the increased relative solubility of the proteins in the surrounding solvent (Supplementary Figure 2A). The condensates also appeared to maintain a constant degree of overlapping area and demonstrated coupled displacement throughout the trajectory. Their overlapping area oscillated around an average value of 40% (Supplemental Figure 2B), which was about 33% lower than the average overlap for α = 0.98. This decrease was sensible given the expansion and irregular shape of the condensate with weaker protein-protein interactions. Despite the decreased overlapping area, we observed strong displacement alignment (Supplemental Figure 2C). Compared to the α = 0.98 condition, we observed a slight rightward shift in the distribution, with a modest increase from 50% to 55% of displacements occurring above 0.6.

When the α = 1.00 scaling factor was applied, the condensates appeared more rounded and compact (Figure 3A) compared to those with α = 0.98 scaling (Figure 1A). The upper and lower leaflet condensates maintained overlapping area and appeared to diffuse together until 10 µs (Figure 3A, panels 1-3). At this point, the two condensates gradually separated until no overlap was observed (Figure 3A, panels 3-6). Concurrently, the overlapping area gradually decreased midway through the trajectory until reaching zero at the 17 µs timepoint (Figure 3B). Analysis of the displacement alignments revealed a leftward shift in the distribution compared to the α = 0.98 condition (Figure 3C). Whereas most displacements occurred with an alignment above 0.6 with α = 0.98, the majority of displacement now occurred with alignments below 0.6, with the number of completely antiparallel displacements doubling. Only 25% of displacements populated the upper two bins, a 30% decrease from the α = 0.98 scaling condition. Overall, these results demonstrated that coupling responded to the strength of protein-protein interactions in the condensates. Specifically, increasing protein-protein interaction strength inhibited transbilayer coupling.

### Suppression of membrane curvature and fluctuations rescues stable coupling when protein-protein interactions are increased

We then sought to investigate why increasing protein-protein interactions inhibited transbilayer condensate coupling. In order to do this, we looked at how the membrane underneath the condensates changed between α = 0.98 and α = 1.00. Specifically, we measured the lipid height in each bilayer leaflet as an indicator of curvature. In order to represent curvature directly underneath the condensate, we first centered the box around the upper condensate center of mass before dividing the membrane into two-dimensional bins. The height of the headgroups in each bin was then averaged across the last 5 µs of the trajectory and mapped with Gaussian smoothing to represent the average vertical positioning of lipids underneath the protein.

In the bilayer height map with α = 0.98 scaling, we observed a height variation of 0.7nm between the highest and lowest lipid heights (Supplemental Figure 3A). There was a patch of lower average height in the region covered by the upper condensate (brown) indicating a negative curvature relative to the proteins. Additionally, there was a patch of higher average height above the region covered by the lower condensate (darker teal), also indicating a negative curvature.

In contrast, when α =1.00, we observed an average positive curvature towards the condensate mass, which was complemented on the opposing leaflet (Figure 4A). The dark teal region indicates positive curvature and corresponds to the region covered by the centrally located upper condensate. The dark brown region indicates negative curvature and corresponds to the average relative position of the lower condensate. The total range of lipid heights was ∼1.2 nm, or about a quarter of the total bilayer thickness (4-5 nm), and a 1.7x increase over the α = 0.98 condition. Based on these results, it appeared that each condensate induced positive curvature toward itself, inverse to the curvature generated by the opposing condensate, such that both curvatures could not be maintained while preserving overlap. We hypothesized that the competing curvature of the condensates led to the eventual decoupling observed in Figure 3A. To test this hypothesis, we introduced tension to the bilayer, which suppressed transverse membrane fluctuations. Tension was induced by coupling pressure in the xy plane to a −10 bar negative pressure. Beginning with a configuration in which the condensates were partially overlapping, we applied negative pressure coupling in the xy plane. In doing so, we effectively imposed tension on the system and suppressed transverse membrane fluctuations and membrane curvature. We performed an equilibration followed by a 20 µs production simulation and analyzed the membrane curvature following the same protocol as for the no tension condition.

**Figure 4.**
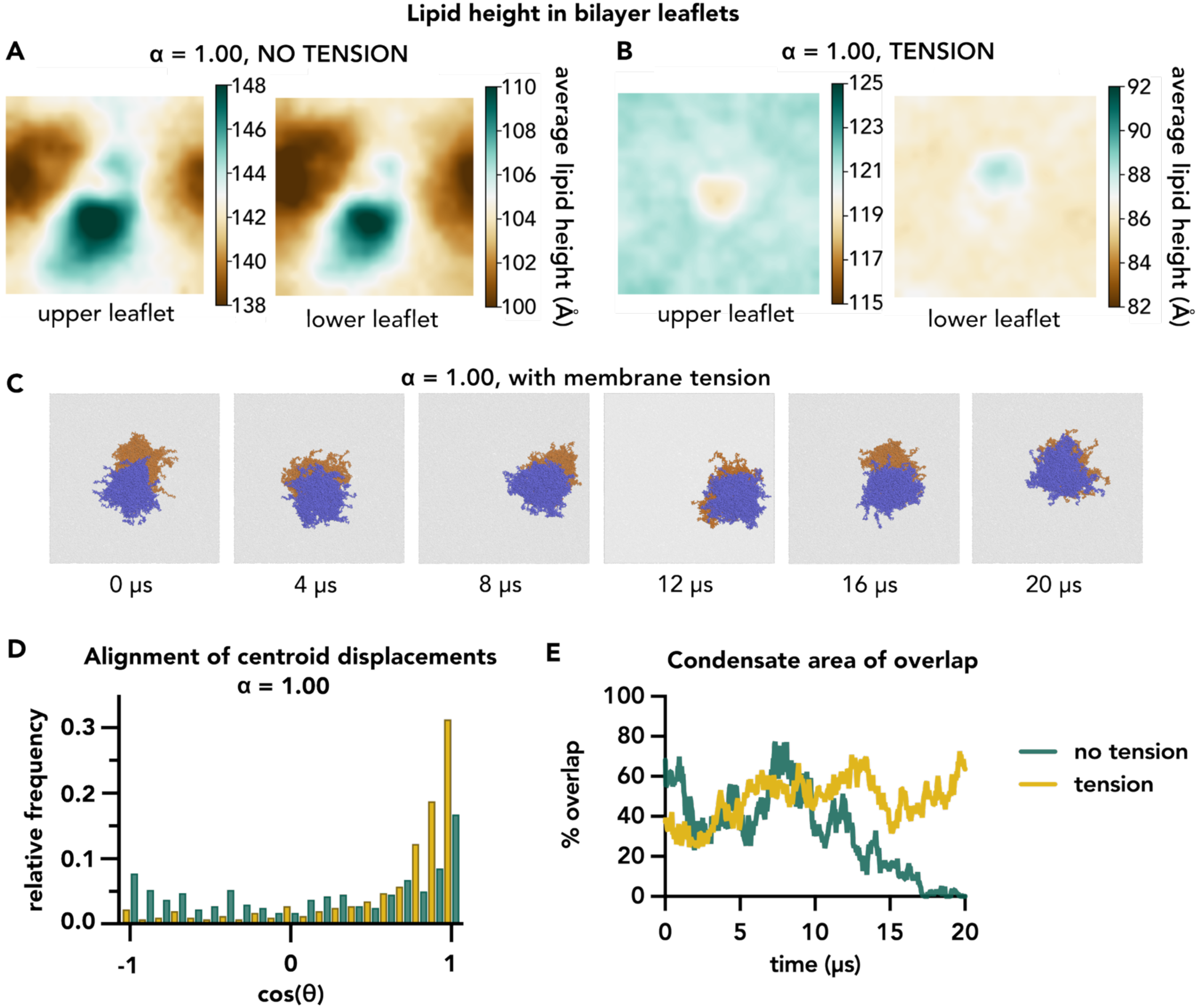
Suppression of membrane curvature and fluctuations rescues full coupling in α = 1.00 scaling condition. **A.** Heatmap of the average height of a PO4 bead in the membrane for a doubled-sided system with α = 1.00 scaling (see Figure 3). The system was centered around the upper leaflet condensate. The contrast range was set to the 10 Å to match the range of the data and centered around the data median. A region of relatively increased average height (dark teal) was observed in both leaflets and was slightly offset from the position of the upper-leaflet condensate. A region of reduced height (dark brown) also appeared in both leaflets, roughly corresponding to the position of the lower-leaflet condensate. **B.** Heatmap of the average PO4 bead height for a double-sided system with α = 1.00 scaling and tension applied in the xy plane, centered on the upper-leaflet condensate. For ease of comparison, the contrast range matches that in A, with the midpoint adjusted to the trajectory median. A region of relatively decreased average height occurred in the center of the upper leaflet (indentation from upper leaflet surface toward the central plane of the bilayer), corresponding to the area covered by the upper condensate. A region of relatively increased average height occurred in the center of the lower leaflet (indentation from lower leaflet surface toward the central plane of the bilayer), corresponding to the area covered by the lower condensate. **C.** Time series snapshots from a trajectory with α = 1.00 scaling and tension applied in the xy plane, viewed from above the plane. The upper condensate (blue) and lower condensate (orange) maintained overlap for the duration of the trajectory and appeared to diffuse together. **D.** Distribution of the alignment of in-plane (xy) displacements of the upper and lower leaflet condensate centroids for α = 1.00 scaling with tension (yellow, C) in comparison to α = 1.00 without tension (green, Figure 3C). **E.** Comparison of overlapping condensate area between α = 1.00 scaling with tension (yellow) and without tension (green, Figure 3B). Overlap was maintained for the duration of the trajectory when tension was applied but decreased to zero without tension.

The map of the average lipid height in the bilayer revealed a much lower range of lipid heights once tension was applied (Figure 4B). In fact, we observed small regions of negative indentation underneath the condensates in each leaflet an order of magnitude lower than in previously measured (Figure 4A). Notably, this reduced curvature coincided with a “rescue” of coupled diffusion of condensates on opposite sides of the membrane (Figure 4C-E). The condensates maintained overlap and co-diffused for the duration of the 20 µs trajectory (Figure 4C). Additional analysis of displacement alignment (Figure 4D) demonstrated a similar distribution as the α = 0.98 and 0.9775 scaling conditions (figures 1C and 2C). 50% of displacements had an alignment of over 0.85, populating the upper two bins, while 68% were aligned greater than 0.6, a 36% increase from α = 0.98. The overlapping area of the system with tension applied gradually increased until converging after 5 µs to an average of ∼34% (Figure 4E), in contrast to the system without tension where overlap decreased to zero. Taken together, these results suggested that curvature acted to inhibit coupling, while dampening membrane curvature rescued the condensates’ ability to couple across the bilayer.

To determine whether the induced curvature was specifically a result of having two condensates on opposing leaflets or if curvature could be induced in a single-sided system, we also simulated a system with a condensate on only one leaflet for both α = 0.98 and α = 1.00 scaling factors (Supplementary Figure 3). Additional simulation details are provided in methods and SI. We included only the final 5 µs of the trajectories in our analysis. In the α = 0.98 condition, the lipid height was higher underneath the condensate in both leaflets, indicated by the dark teal region in the membrane center (Supplemental Figure 3C). The difference between the highest average point and lowest average point was ∼1.2 nm. Similarly, when α = 1.00, the average lipid height was higher underneath the condensate in both leaflets, indicating positive curvature towards the proteins (Supplemental Figure 3D). However, the range was reduced by ∼20% from the α = 0.98 single sided system to 1 nm. In both conditions, the induced curvature was not dependent on the presence of a condensate on the opposite leaflet.

### Entropy was reduced for lipids in direct contact with the proteins but there was no effect on the opposing leaflet

Given the apparent stability of the coupling, we next investigated the factors contributing to the underlying energy landscape. Previous work has proposed that condensate coupling is driven by an entropic force which favors consolidation of regions with reduced entropy that arise from condensate-induced reduction of lipid mobility.^26^ In the uncoupled state, a population of lipids located opposite the protein condensate (Figure 5B, “mirrored”) was proposed to exhibit reduced entropy via interleaflet interaction. Coupling of the condensates (Figure 5A) is therefore hypothesized to increase the total entropy of the system by relieving the entropic constraint on the two mirrored regions. If so, we expected that the entropy of a coupled system should maximize the system entropy such that

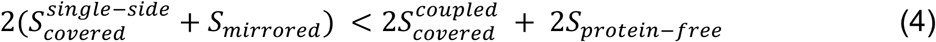

where S is the entropy of a single lipid in the regions defined in Figures 5A and 5B. For regions of equal size and containing the same number of lipids, the term corresponding to the number of lipids may be omitted. Accordingly, the following analysis compared per-lipid entropy values rather than total entropy summed over the total regions.

**Figure 5.**
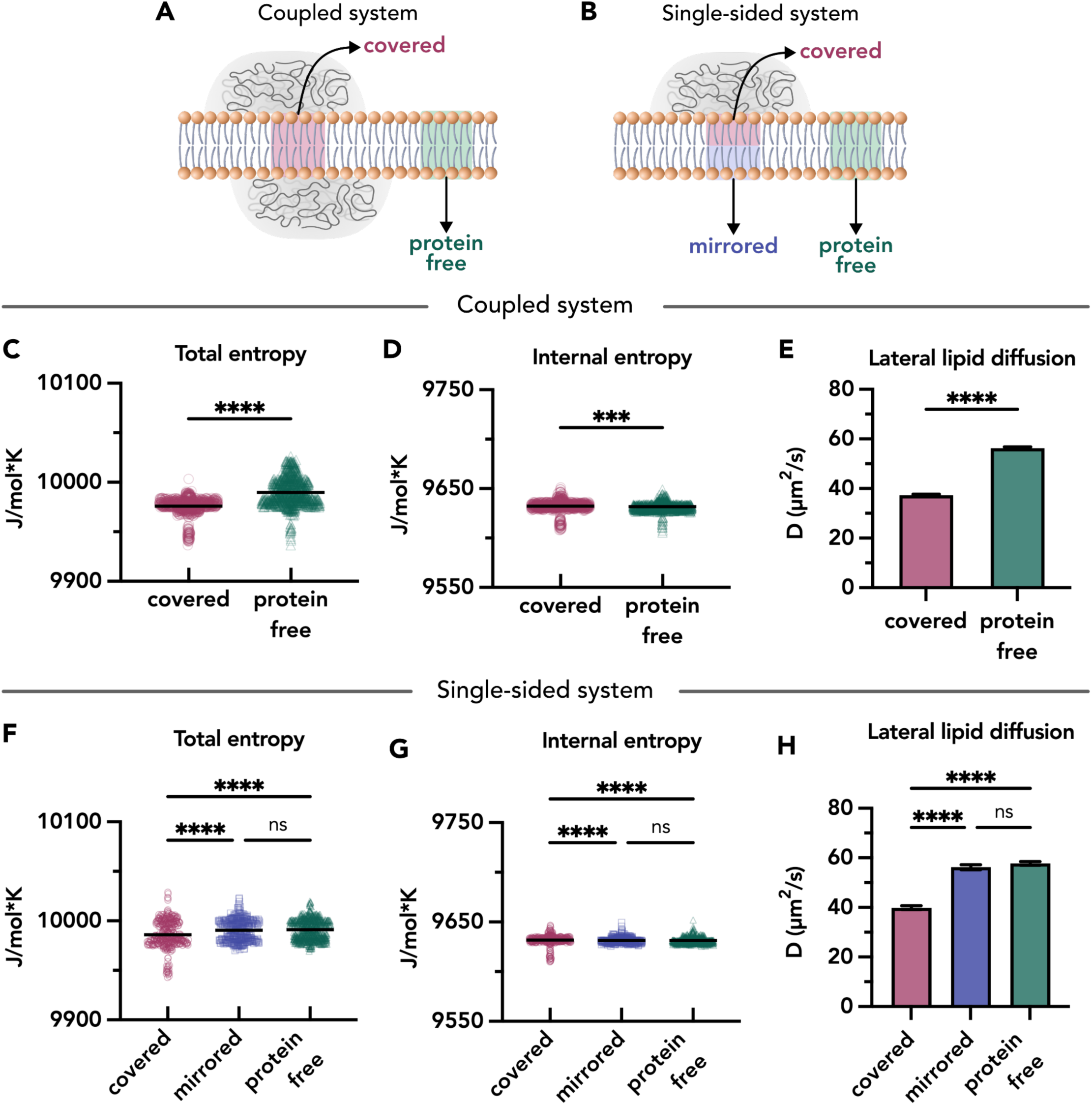
Entropy was reduced for lipids in direct contact with the proteins but there was no effect on the opposing leaflet. **A.** Illustration of lipids in a coupled system. Lipids directly underneath and in contact with either the upper leaflet or the lower leaflet condensate are labeled “covered”. Lipids which are not in contact with condensate are labeled “protein free”. Proteins are rendered grey. **B.** Illustration of lipids in a single-sided condensate system. Lipids which are directly opposite to the condensate but in the opposing leaflet are labeled “mirrored”. Covered and protein free lipids are defined as in A. **C.** Total quasi-harmonic entropy calculated for both covered (pink) and protein free (green) lipids in a coupled system, with coloring corresponding to the groups labeled in A. Protein free lipids demonstrated higher total entropy compared to covered lipids. Data was sampled from the production simulation shown in figure 1. Symbols represent the entropy of a single lipid and bars mark the sample mean. **D.** Internal quasi-harmonic entropy calculated for lipids in C. Covered lipids had a higher measured internal entropy compared to protein free lipids. Translational and rotational fitting was performed prior to calculation of entropy. **E.** Lateral lipid diffusion of covered and protein free lipids in a coupled system. Covered lipids had a lower measured diffusion coefficient than protein free lipids. Data was sampled from all three repeat simulations. Bars show the mean of individual lipids with standard error. **F.** Total quasi-harmonic entropy calculated for covered (pink), mirrored (blue) and protein free (green) lipids in a single-sided system, with coloring corresponding to the groups labeled in B. The total measured entropy was lowest for covered lipids compared to both mirrored and protein free lipids. Mirrored and protein free lipids did not demonstrate a measurable difference in total entropy. Data was sampled from a single production simulation. **G.** Internal quasi-harmonic entropy calculated for lipids in F. There was no measured difference in the internal entropy of the three groups. Translational and rotational fitting was performed prior to entropy calculation. **H.** Lateral lipid diffusion constants for covered (pink), mirrored (blue) and protein free (green) lipids in a single-sided system. The measured diffusion constant was lowest for covered lipids in comparison to both mirrored and protein free lipids. Mirrored and protein free lipids diffusion constants were not statistically distinguishable. Data was sampled from all three repeat simulations. Bars show the mean of individual lipids with standard error. *Coupled system*: Mann-Whitney non-parametric comparison tests (***p < 0.001, ****p < 0.0001), *Single-sided system*: Kruskal-Wallis with uncorrected Dunn’s multiple comparison tests (****p < 0.0001)

We quantified lipid entropy using the quasi-harmonic approximation proposed by Schlitter, an approximate yet widely accepted method that yields results from coarse grained simulations consistent with those obtained from more detailed atomistic approaches^36,37^. The total entropy of a molecule (Figure 5C & 5F) is computed from its raw coordinates and captures contributions from all modes of motion. The internal entropy is obtained by removing translational and rotational motion through fitting to the molecular center of mass. For lipids, the internal entropy reflects bond rotation and stretching, and encapsulates acyl chain ordering. A detailed description of entropy calculations is described in methods.

We first examined the lipid entropy in a coupled system (Figure 5A). For each analyzed trajectory frame, we selected lipids within a 4 nm radius of the centroid of both condensates, excluding dilute phase chains. These lipids were labeled as “covered” (Figure 5A, pink). We also made a 4 nm radius selection of lipids from a region of the membrane which was unassociated with either condensate, labeled “protein-free” (Figure 5A, green). We next evaluated the entropy of individual lipids within the selected regions (Figures 5C and 5D). First, analysis of total entropy revealed that lipids in protein-covered regions exhibited an average entropy of 9976 J/mol*K (Figure 5C, pink), compared to 9990 J/mol*K for protein-free lipids (Figure 5C, green), representing a modest decrease of 14 J/mol*K. In addition, the protein-free population showed ∼60% greater variance (13.6 vs. 8.4 J/mol*K), further supporting a distinction between the two populations. Next, we performed translational and rotational fitting on the same lipid populations analyzed in Figure 5C to isolate their internal entropy (Figure 5D). In contrast to the total entropy, we found that although the two lipid populations were statistically distinguishable, their mean internal entropies differed by less than 1 J/mol*K, with the covered lipids (Figure 5D, pink) exhibiting a slightly higher mean than the protein-free lipids (Figure 5D, green). Additionally, the distributions were also visually similar in shape and their variances differed by only 1.5 J/mol*K. We therefore concluded that the apparent statistical significance arose primarily from the large sample size (n > 900 per group), rather than a meaningful physical difference. These results suggested no significant change in acyl chain ordering under the coupled condensates. Overall, we observed reduced total entropy for covered lipids compared to protein free lipids, but no differences in internal lipid entropy, where the translation and rotational movement were removed. We therefore hypothesized that reduced translational and rotational motion were the primary contributors of entropy reduction for covered lipids in the coupled system.

To further quantify the differences between the condensate-covered and protein-free, we next calculated the lateral diffusion constant for the two groups. We found that covered lipids had an average lateral diffusion constant of 37 µm^2^/s compared to an average of 56 µm^2^/s for lipids in a protein-free region, a ∼30% reduction (Figure 5E, pink vs. green). Lower measured diffusivity of covered lipids is consistent with the conclusion made from the quasi-harmonic entropy calculations that the condensate reduces local lipid entropy.

Having demonstrated an entropy reduction in the protein-covered region of the coupled system, we next asked whether the entropy reduction was transmitted through the bilayer and if the coupled system was more entropically favorable than a system of two non-overlapping condensates. To enable the comparison described in Equation 4, we repeated our analysis of entropy for lipids in a single-sided condensate system as an analog for an uncoupled condensate (Figure 5B). For each analyzed trajectory frame, we selected lipids within a 4 nm radius of the condensate centroid, and split into covered lipids, which were directly underneath the condensate (Figure 5B, pink) and mirrored lipids which were in the same region but in the opposite leaflet (Figure 5B, blue). We again made a radius selection of protein free lipids from a region of the membrane which was unassociated with the condensate (Figure 5B, green).

First, we evaluated the total entropy of lipids in each group. As a sanity check, protein-free lipids exhibited the same average entropy of 9990 J/mol*K in both the coupled and single-sided systems (Figure 5F vs. Figure 5C, green). In contrast, the average total entropy of covered lipids in the single-sided system was 5.1 J/molK lower than that of protein-free lipids (Figure 5F, pink), approximately one-third of the difference observed between these groups in the coupled system. Mirrored lipids showed a similarly higher average total entropy than covered lipids by 4.5 J/mol*K (Figure 5F, blue). Notably, we did not observe a statistically significant or substantial difference in total entropy between mirrored and protein-free lipids. Analysis of the internal entropy revealed trends similar to those observed in the coupled system (Figure 5G). Although statistically significant differences were detected between covered and protein-free lipids and between covered and mirrored lipids, these differences were less than 1 J/mol*K and therefore not substantial relative to the total entropy, suggesting no meaningful physical differences between the populations.

Investigation of lateral lipid diffusivity reinforced these findings (Figure 5H). While no difference in diffusion coefficient was observed between the mirrored and protein-free groups, covered lipids exhibited an approximately 30% reduction in diffusion coefficient relative to protein-free lipids. Compared to the coupled system, covered lipids in the single-sided system displayed an increased diffusion coefficient of 2.58 µm^2^/s, or 7%. Taken together with the corresponding increase in total entropy, this result suggested that condensate coupling imposed a small but measurable restrictive effect on lipid mobility in the covered region.

Overall, these data did not support the hypothesis of an entropic loss for mirrored lipids that could be recovered through coupling and proposed in equation 4. Instead, the simulations suggest that because the entropy of mirrored and protein free lipids was indistinguishable, lipid entropy was not the primary driving mechanism for stable condensate coupling in our system.

### RGG condensates induce lipid tail interdigitation in the protein-covered region

We did not identify an entropic mechanism for coupling in our system. However, more broadly, multiple mechanisms have been proposed for coupling between lipid domains across a bilayer, including chain interdigitation and local suppression of membrane fluctuations^38^. We therefore sought to determine whether any of these mechanisms were present in the coupled condensate system.

We first investigated the lateral density of lipids in the protein-covered and protein-free regions (Figure 6A), as described in the Methods. The average lateral density in the protein-free regions was 1.73 lipids/nm^2^, denser in comparison to literature values for POPC bilayers which range from 1.43-1.58 lipids/nm^2^ ^39–42^. However, when calculated over the entire simulated membrane, the lateral density was 1.55 lipids/nm^2^, consistent with reported values. This discrepancy suggested that the analysis performed on the sampled regions was subject to a methodological error. Regardless, the density values we calculated were informative relative to each other. We found that lipid density was depleted by 4% in the covered regions (Figure 6A, pink) compared to the protein-free region (Figure 6A, green) in our system.

**Figure 6.**
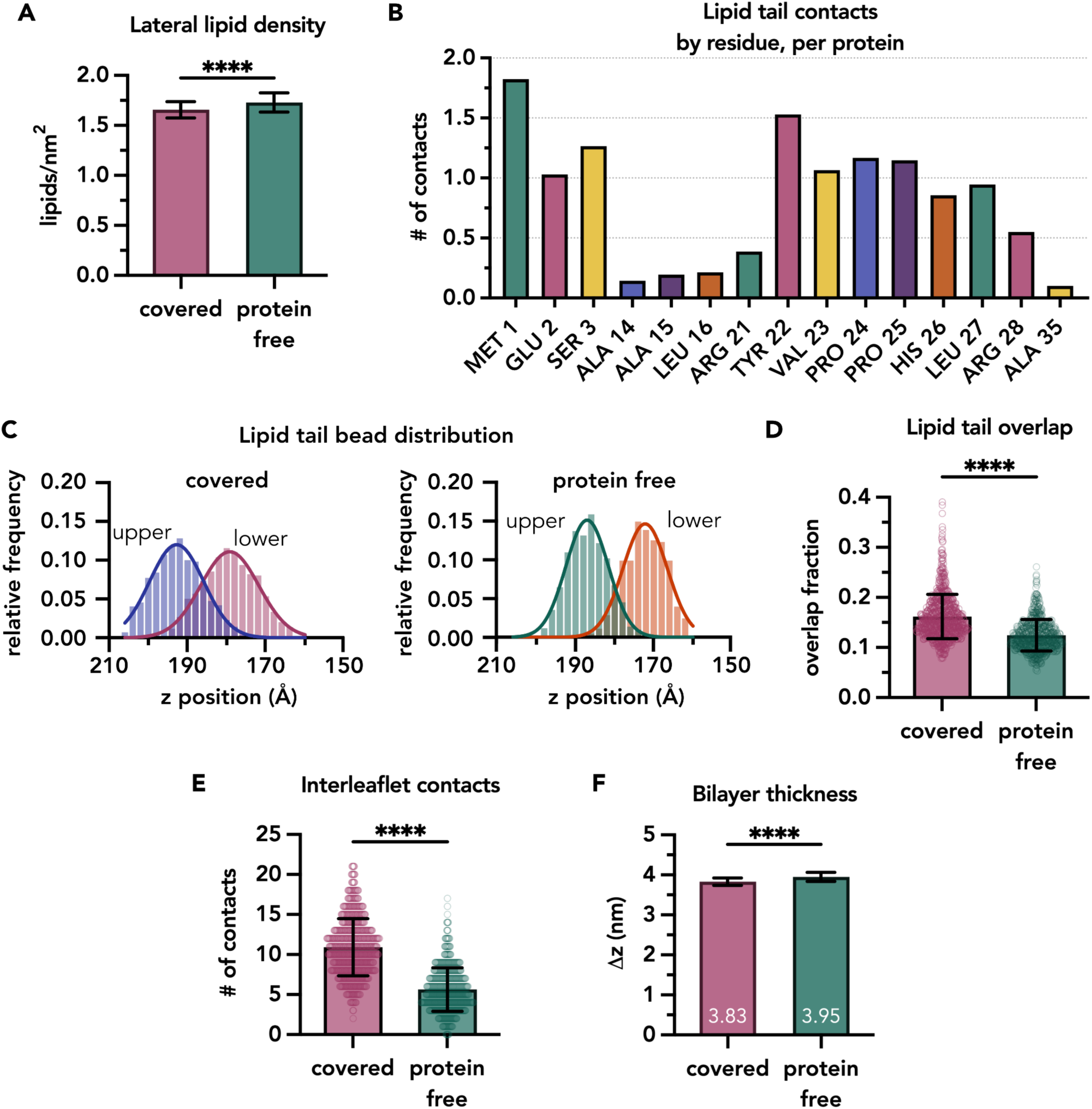
RGG condensates induce lipid tail interdigitation in the protein-covered region. **A.** Lateral lipid density of protein-covered and protein-free lipids averaged across three trajectories. Data was sampled every 50,000 ps from three repeat trajectories. Error bars show std dev. Unpaired t test with Welch’s correction (****p < 0.0001). **B.** Number of contacts between lipid tail acyl beads and protein residues for the top 15 residues. Contacts were defined using a 5 Å cutoff. Values were summed over 200 frames and normalized by both the number of frames and the total number of individual proteins. For example, in each protein chain, residue *X* is in contact with *y* acyl beads. Residues MET 1 and GLU 2 are tethered to a lipid headgroup. **C.** Distributions of lipid tail acyl bead z-positions in the membrane for a protein-covered region (left) and a protein-free region (right), overlaid with Gaussian fits. Blue and pink data represent the upper and lower leaflet distributions, respectively, in the protein-covered region, while green and orange curves represent the upper and lower leaflets in the protein-free region. These example distributions are representative of the summary data shown in D. **D.** Degree of lipid interdigitation in protein-covered (pink) and protein-free (green) regions, quantified as the fractional overlap of Gaussian distributions describing lipid tail z-positions of each leaflet. The overlap corresponds to the shared area between the two Gaussian distributions, as illustrated in C. Overlap was sampled every 50,000 ps across three repeat trajectories. Each symbol represents a single frame. Error bars show std dev. Mann-Whitney non-parametric comparison test (****p < 0.0001). **E.** Number of contacts between upper and lower leaflets. Contacts were defined as between lipid tail beads from opposing leaflets within a 5 Å cutoff distance. Data was sampled every 50,000 ps from three repeat trajectories and averaged per frame. Each symbol represents a single frame. Error bars show std dev. Unpaired t test with Welch’s correction (****p < 0.0001). **F.** Bilayer thickness in sampled regions for protein-covered (pink, left) and protein-free (green, right) regions. Thickness in a region was calculated by averaging the height of all headgroup beads in the upper and lower bilayer leaflets and taking the difference. Data was sampled every 50,000 ps from three repeat trajectories and averaged per frame. Error bars show std dev. Unpaired t test with Welch’s correction (****p < 0.0001).

We hypothesized that this local depletion arose from the steric bulk of the protein condensates at the membrane surface or from residue insertion into the bilayer. To test this, we quantified the extent of residue insertion beyond the lipid headgroup region by measuring contacts within 5 Å between protein residues and lipid tail acyl beads. Contacts were summed over the course of the trajectory for each residue and then averaged per frame. This per-frame average was subsequently normalized by the total number of proteins in the system and reported for the top 15 residues (Figure 6B). The resulting values can be interpreted such that, for a given protein chain, *residue X* is in contact with *y* acyl beads.

Analysis of protein-lipid tail contacts showed that the highest number of contacts occurred for the first three residues as a group. This result was expected due to the constant force tether between the terminal residues and a lipid headgroup. These residues penetrated into the bilayer, with each residue forming at least one acyl bead contact in each protein chain on average. Among the remaining top residues, all but two were hydrophobic, suggesting that insertion occurred through non-specific hydrophobic interactions, consistent with reports of residue positing in lipid bilayers, including for intrinsically disordered proteins^43–46^.

Although we measured a slight decrease in lateral lipid density beneath the condensates and identified a mechanism underlying this effect in our system, our observations were not consistent with previous literature which describes lipid packing behavior induced by biomolecular condensates. This prior study of membrane wetting by glycinin condensates reports increased local lateral lipid packing, inferred from a spectral shift in a membrane-embedded dye^47^. However, the system differs from ours in membrane composition, condensate composition, and the attachment between the condensate and membrane. It therefore may exhibit distinct behavior. Even so, we considered alternative mechanisms that could reconcile our observations. An increase in area per lipid, or a decrease in lateral headgroup density, has been associated with increased acyl chain interdigitation^48–51^. Given the apparent increase in per lipid area in our system resulting from protein-lipid tethering and residue insertion, we next investigated the extent of lipid tail interdigitation in the coupled system.

We quantified the degree of lipid tail interdigitation (Figure 6C and 6D) using similar 2nm radius cylindrical selections as in previous analyses. The limited area sampled was intended to minimize variance due to effects from spontaneous curvature. To obtain the overlap fraction, we separated the lipids into top and bottom leaflets and obtained the z positions of the acyl tail beads. We then fit the distributions of each leaflet to a Gaussian function and integrated the area of overlap between them to obtain an overlap fraction (Figure 6C and 6D). The sample distributions in Figure 6C are representative of the average overlap values for covered and protein free lipids which are plotted in Figure 6D. Generally, we observed that lipid tail height distributions in the protein covered region (Figure 6C, left) appeared broader in comparison to those in the protein-free region (Figure 6C, right). Protein-free lipids had an average overlap fraction of 0.122 (Figure 6D, left, green) while covered lipids had an average overlap of 0.163 (Figure 6D, left, pink), a 34% increase. In order to confirm that our observations were not exclusively a result of geometry fluctuations or methodological error, we additionally quantified the number of interleaflet contacts between lipid acyl tail beads, which was agnostic to variations in membrane height (Figure 6E). Supporting our observations of leaflet overlap, the average number of contacts between the upper and lower bilayer leaflets in the covered region was doubled compared to a protein free region. An increase in tail interdigitation should logically also occur with a reduction in overall membrane thickness. Indeed, when we quantified the membrane thickness, we found that the covered regions were less thick than the protein-free regions by about 1 Å, or ∼2.5 % of the total membrane thickness in the protein free region.

To further investigate the degree to which interdigitation is due to the condensate-bilayer interaction or specifically coupling of the condensates, we then repeated each of these analyses for the single sided system (Supplemental Figure 4A-4C). We observed a similar trend in lipid tail overlap between the covered and protein free regions in the single-sided system. Covered regions had an average leaflet overlap fraction of 0.153 compared to 0.123 for protein free regions, a 24% increase (Supplemental Figure 4A, hash bars). While the protein-free regions in the coupled and single-side systems were indistinguishable, the average overlap fraction in the covered region was 6% higher in the coupled system than the single-sided system. We also found that the average number of interleaflet contacts was lower on average by 0.5 and the bilayer thickness was increased by 0.4 Å in the covered region of the single-sided system (Supplemental Figure 4B and 4C, respectively). Although these differences were small relative to the magnitude of the measurements, they were statistically significant and suggested a compounded effect on the underlying lipids when condensates were present on both leaflets.

Overall, the data suggests that condensate residues inserted into the bilayer, leading to a local decrease in lateral lipid density. The resulting increase in per lipid area then promoted increased interdigitation between leaflets relative to protein free regions. Based on these observations, we hypothesized that when two condensates overlapped, interdigitation between the covered leaflets resisted their separation, thereby stabilizing the coupled behavior.

## Conclusions

Protein condensates are increasingly understood as drivers of membrane organization in numerous biological contexts. Notably, they are capable of inducing lipid domains and are essential in processes in which protein assemblies are coordinated across the membrane. Recent work demonstrates that IDP condensates are stably coupled when allowed to form on opposite leaflets. That is, without any membrane spanning domains, two opposing condensates are able to communicate through the bilayer. However, the mechanisms underlying transbilayer communication are not evident. In order to probe the molecular behavior underlying this phenomenon we performed coarse grain molecular dynamics to simulate a system of condensates on opposite sides of a lipid bilayer under various conditions.

We placed two pre-formed condensates of RGG on the surface of each bilayer leaflet, such that their projections in the xy plane were overlapped. Using a value from previous work, we applied an α = 0.98 to scale down all protein-protein interactions. We showed that mirrored condensates in the simulation are stably coupled during simulations. However, increasing the strength of protein-protein interactions (α = 1.00) led to the decoupling of the condensates. We found that in this condition positive membrane curvature was increased underneath each condensate and these opposing curvatures likely inhibited coupling. We then applied tension to the bilayer to suppress height fluctuations and showed that removing curvature resulted in the rescue of stable coupling even with the increased protein-protein interactions.

Next, we examined the lipids in the system to understand the mechanism of transmembrane coupling. We examined the lipids which were directly covered by the protein condensates and found that their total entropy was reduced relative to protein-free lipids. However, when translational and rotational motion were removed, the two populations were indistinguishable, indicating that the entropy reduction primarily arose from decreased lipid mobility. Consistent with these results, covered lipids exhibited reduced lateral diffusion compared to protein free lipids. Notably, when a condensate was present on only one leaflet, lipids in the opposing leaflet did not show reduced entropy or diffusivity. These findings demonstrated that entropy reduction was not transferred across the bilayer, and not responsible for driving condensate coupling in our system. In contrast, we observed that while lateral lipid packing decreased in condensate-covered regions, lipid tail interdigitation (transverse packing) increased significantly relative to protein-free regions. Together, these results support a model in which enhanced interleaflet overlap serves as the primary mechanism of information transfer and the dominant stabilizing factor for transmembrane condensate coupling.

The observed differences in lipid tail interdigitation and leaflet overlap between condensate-covered regions in the coupled and single-sided systems were statistically significant but small relative to the overall measured values of lipid tail overlap, interleaflet contacts, and bilayer thickness. These small differences likely reflected the limited resolution of the model, in which each coarse-grained bead represented 3-4 carbon atoms and bond lengths were ∼0.5 nm. Variations below this scale are likely approaching the limit of detection of the model. Higher-resolution approaches, such as atomistic simulations, would likely be required to more precisely quantify these subtle but potentially important differences in lipid interdigitation.

In addition, due to computational limitations on the system size, we may not observe effects that emerge only in macroscopic systems, such as large scale membrane fluctuations. For example, we observed no measurable changes in lipid entropy that would result from coupling two condensates across the membrane in our simulation systems (Figure 5). Our condensates are 10 nm diameter, several orders of magnitude smaller than experimentally measured micron-scale condensates which may have a stronger effect on lipid order and fluctuations in both leaflets. Future studies will build on this work by employing atomistic models to resolve subtle effects and more coarse large scale models to capture contributions from membrane fluctuations.

## Acknowledgements

PR thanks the support from the Welch Foundation under F-2021. JCS acknowledges support from the National Institutes of Health under R35GM139531, including a supplement to support KZ, and from the Welch Foundation under F-2257. PR and JCS additionally thank support from NSF BIO under 2529782. KZ further acknowledges a graduate research fellowship from the National Science Foundation.

## Author contributions

KZ contributed conceptualization, formal analysis, investigation, validation, methodology, visualization, and writing. PR and JSC contributed conceptualization, formal analysis, investigation, funding acquisition, supervision, and writing.

## Methods

The MARTINI 3.0 force field^28,29^ and GROMACS^52,53^ engine were used for all MD simulations. Structures and trajectories were visualized using pymol^54^. The initial all-atom structure for the protein was taken from an AlphaFold structure prediction of the 170 residue sequence of RGG. This structure was then coarse grained using martinize2^55^ and solvated using insane^30^. Short minimization, equilibration, and production simulations were then performed using the recommended CHARMM-GUI^56,57^ protocols to obtain a more compact conformation for membrane tethering. We assembled a 50 x 50 nm^2^ bilayer using CHARMM-GUI, then performed minimization, equilibration and a short production simulation. After, we removed the solvent and used the dry bilayer as a building block. Then, proteins were added in the desired conformation and the protein-membrane system was solvated with water and 100mM NaCl using *insane*^30^. Simulations were run at 300K in the NPT ensemble with semi-isotropic pressure control coupled to 1 bar in the xy plane and 1 bar along the z axis using the C-rescale barostat, unless otherwise noted.

### Tethering

We attached proteins to the bilayer using a constant force pull-coordinate between the center of mass of the terminal two residues of each chain and the nearest lipid headgroup at the end of minimization. This force was maintained throughout equilibration and production.

### Scaling

The MARTINI force field uses a Lennard-Jones potential to describe non-bonded Van der Waals interactions. In order to control protein-protein interaction strengths, we added a general scaling factor α, to the default parameters:

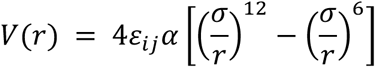

We excluded bead types used to describe lipids, water, and salt ions.

### Tension

In order to impose membrane tension, pressure in the xy plane was coupled to −10 bar, while the z axis remained coupled to 1 bar. Simulation parameter files are available in the SI.

### Calculation of quasi-harmonic lipid entropy

We calculate the quasi-harmonic entropy using the formulation proposed by Schlitter. We selected covered lipids within a 2 nm radius from the in-plane condensate centroid and the protein-free lipids from a 2 nm radius patch in a region which was not covered by protein. For each lipid in the selection, we used *gmx covar* to obtain the covariance matrix and its eigenvalues over a 0.25 µs simulation window from the point of selection. To obtain the internal quasi-harmonic entropy, we included translational and rotational fitting to a reference structure. We then calculate the quasi-harmonic entropy at 300K as

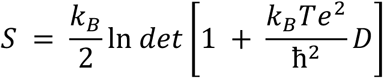

where D the diagonalized covariance matrix, whose determinant is found from the product of the eigenvalues.

### Calculation of diffusion coefficients

Lipid diffusion constants were computed for lipids within a 2 nm radius patch. Trajectories were analyzed over non-overlapping 1 µs intervals centered on the frame at which the selection was made (t_frame_ − 0.5 µs to t_frame_ + 0.5 µs). The first 0.5 µs of each trajectory was omitted. Diffusion coefficients were obtained using the built-in *gmx msd* function and fitting the first quarter of the MSD curve. Analysis was performed for each lipid individually. Outliers were then removed prior to averaging.

Data was sampled every 50,000 ps and averaged per frame from two 10 µs repeat trajectories (single-sided α = 0.98) or one 20 µs trajectory (α = 1.00, α = 1.00 with tension). Symbols represent a single frame when shown. Error bars show std dev.

## Supplementary figures

**Supplementary Figure 1.**
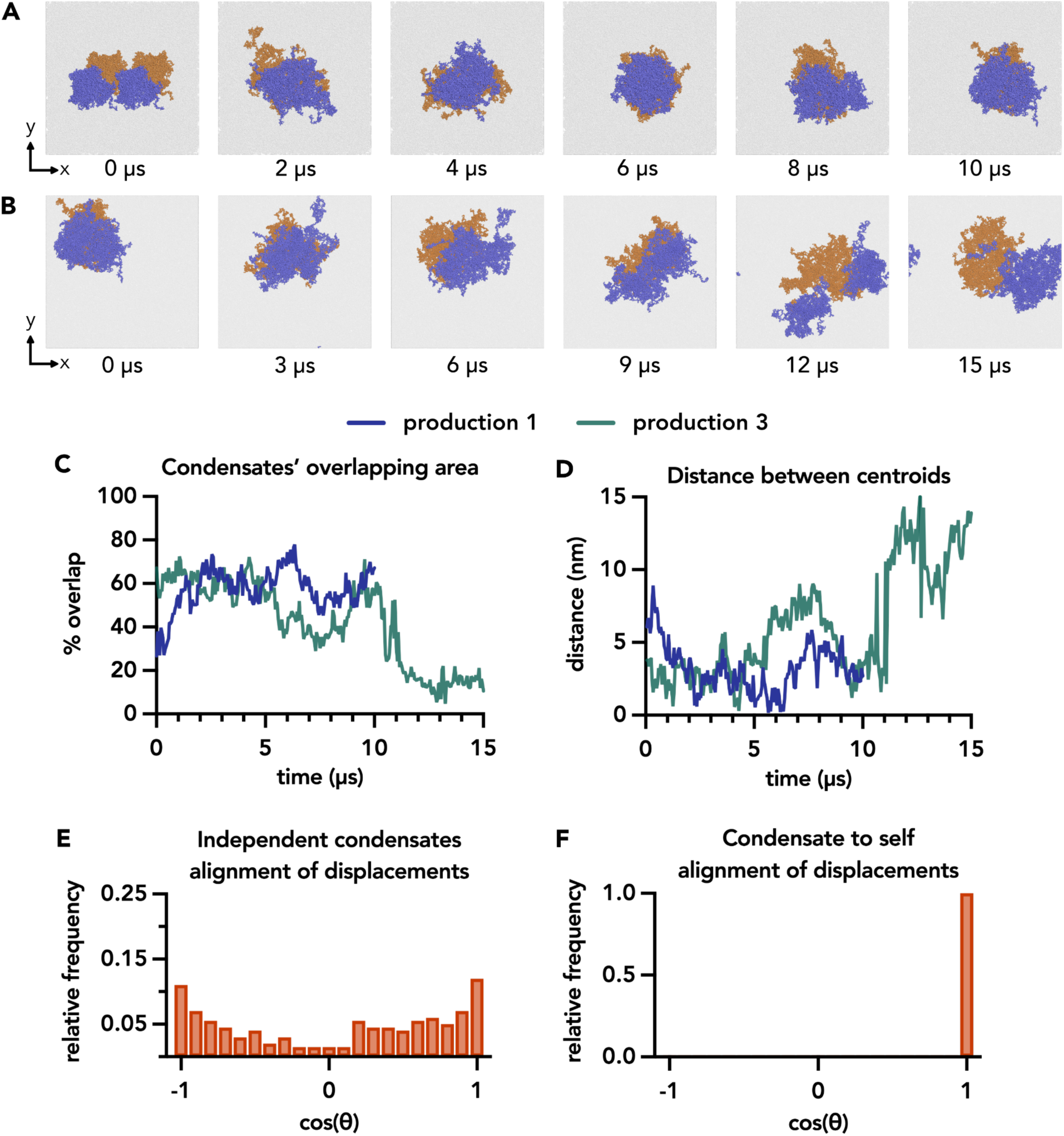
**A.** Representative snapshots of the initial production simulation (production 1) shown from above the xy plane with the bilayer rendered in translucent grey, upper leaflet proteins colored blue, and lower leaflet proteins colored orange. The initial confirmation consisted of two closely placed condensates each containing 16 RGG chains on each leaflet (first panel, 0 µs). These two smaller condensates merged into a single larger condensate containing 32 RGG chains on each leaflet (second panel) within 1 µs. The two condensates maintained overlap throughout the trajectory (panels 2 through 6). The final conformation was used to initialize two additional production simulations (panel 6, 10 µs). **B.** Representative snapshots of an additional repeat trajectory (production 3) shown from above the xy plane with the bilayer rendered in translucent grey, upper leaflet proteins colored blue, and lower leaflet proteins colored orange. The initial position was the same as for production 2. **C.** Overlap of the area projected in the xy (membrane) plane of the condensates shown in A (production 1) and B (production 3), expressed as a percentage of the total condensate area over time (see equation 1). **D.** Distance between condensate centers of mass in the xy plane over the trajectories shown in A (production 1) and B (production 3). **E.** Distribution of the alignment of xy plane displacements of condensates in two iBMndependent simulations. The two unassociated condensates yielded approximately evenly distributed alignments in comparison to the self control, with ∼45% of displacements aligned with a cosine value below zero compared to ∼55% above zero. Displacements for the first condensate were taken from the upper leaflet condensate of the trajectory shown in Figure 1. The second set of displacements was taken from a trajectory of a single sided condensate system. Data was grouped into 21 bins with centers at increments of 0.1 units. Data displayed was taken from alignment over 10 µs, the length of the single-sided system. **F.** Distribution of the alignment of in-plane (xy) displacements of a condensate to itself. The comparison of a condensate to itself yielded exactly parallel alignment with 100% probability, as expected. A trajectory of a condensate from a single-sided system was used. Data was grouped into 21 bins with centers at increments of 0.1 units.

**Supplementary Figure 2.**
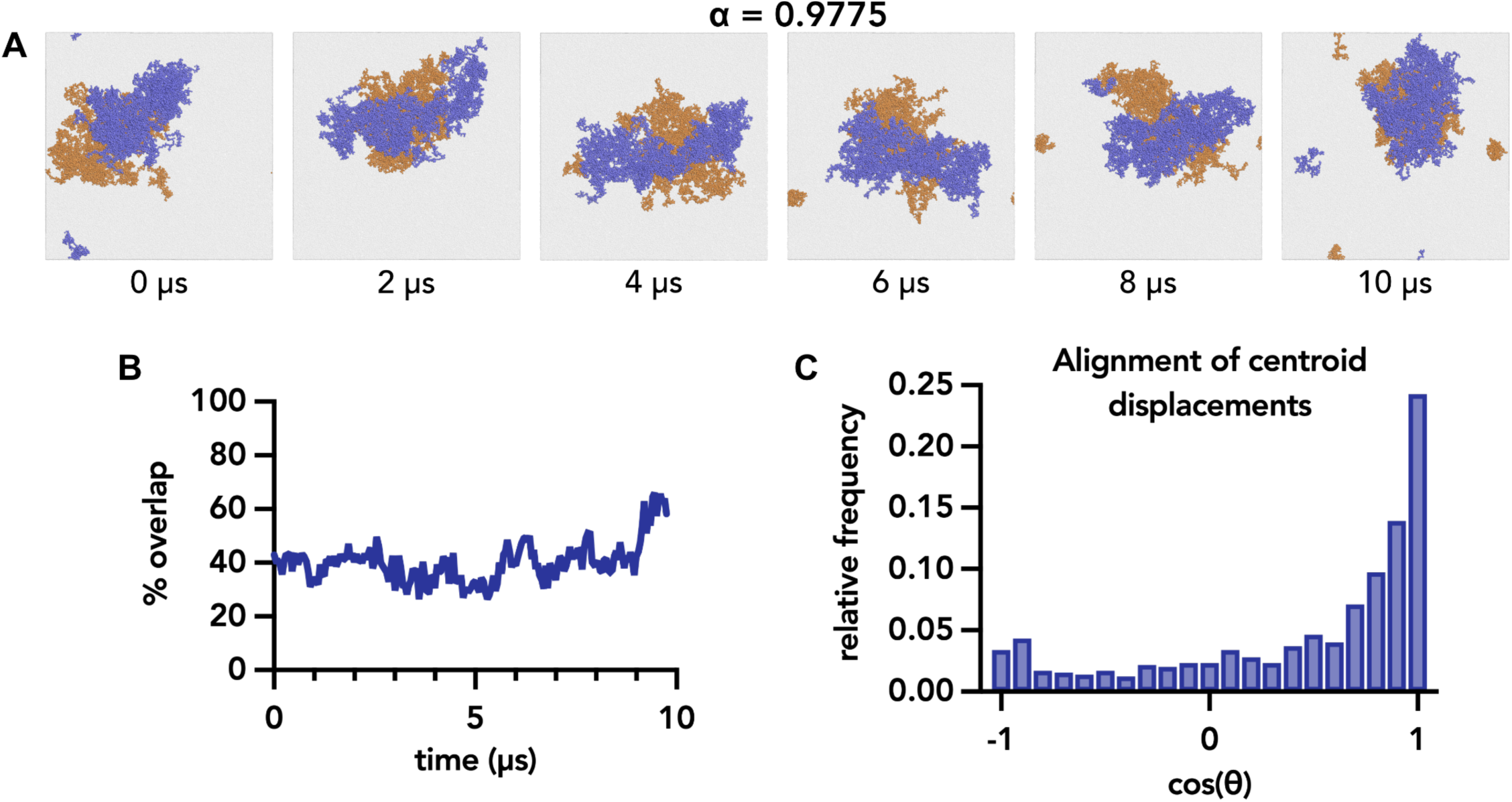
**A.** Representative snapshots from a single repeat shown from above the xy plane with protein-protein VdW interaction strengths decreased by the scaling factor α = 0.9775. Condensates were expanded in comparison to those with stronger protein-protein interactions. The boundaries of the condensates were also less circular and their shapes more irregular. **B.** Overlap of the xy-projected area of the condensates shown in A, expressed as a percentage of the total condensate area over time. The overlapping percentage averaged to ∼40% over the 10 µs shown. **C.** Distribution of the alignment of in-plane (xy) displacements of the upper and lower leaflet condensate centers of mass for three repeat trajectories where α = 0.9775. Condensates diffused with a generally robust alignment. Data was grouped into 21 bins with centers at increments of 0.1 units. Data displayed was taken from three independent trajectories totaling 32 µs.

**Supplementary Figure 3.**
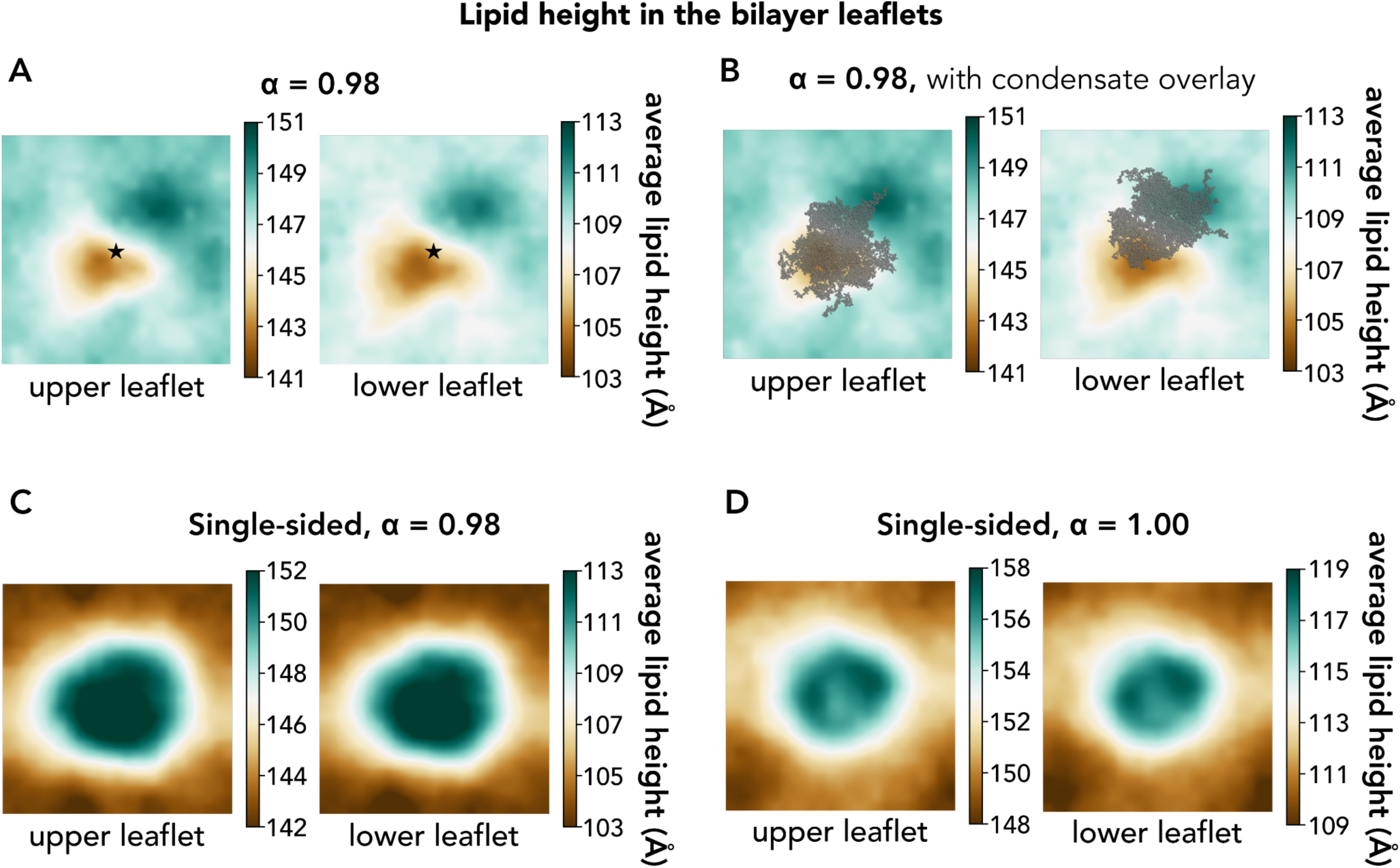
**A.** Map of average height of PO4 lipid headgroup beads in the membrane of the double sided α = 0.98 condition where stable coupling occurred. The membrane center is marked with a black star. The trajectory was centered around the center of mass of the upper condensate. **B.** Plots from A superimposed with representative overlays of condensates to demonstrate the proteins’ positioning on the membrane surface. **C.** Average height of PO4 lipid headgroup beads in a single sided condensates system containing 36 RGG chains with α = 0.98 scaling. **D.** Average height of PO4 lipid headgroup beads in a single sided condensates system containing 36 RGG chains with α = 1.00 scaling.

**Supplementary Figure 4.**
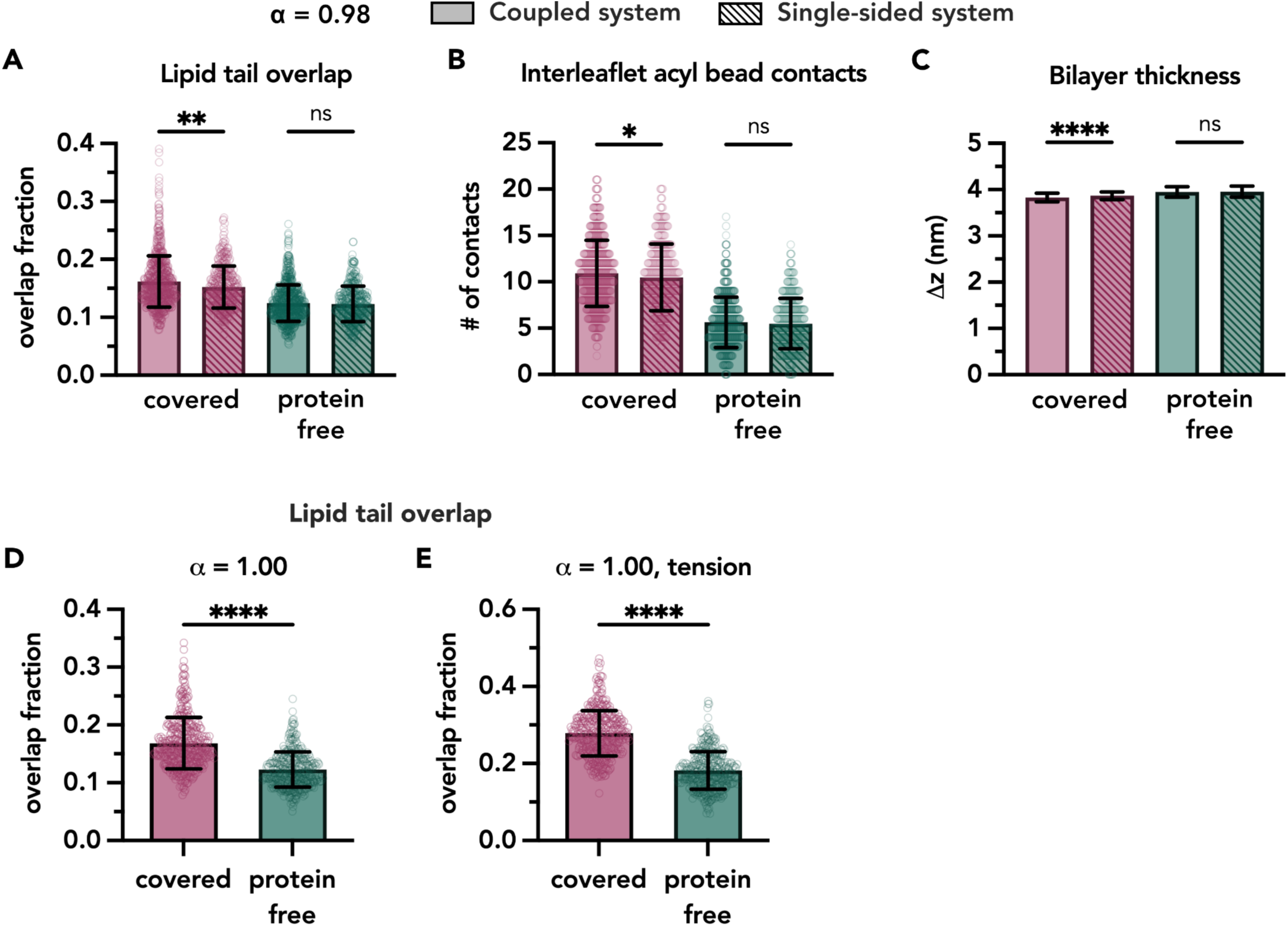
Lipid tail interdigitation in other condensate-bilayer systems. **A.** Degree of lipid interdigitation in the coupled (solid) and single-sided (hashed) condensate systems for protein-covered (pink) and protein-free (green) regions, quantified as the fractional overlap of Gaussian distributions describing lipid tail z-positions of each leaflet. Mann-Whitney non-parametric comparison test (****p < 0.0001). **B.** Number of contacts between upper and lower leaflets with coloring consistent with A. Unpaired t test with Welch’s correction (****p < 0.0001). **C.** Bilayer thickness in sampled regions for coupled vs single with coloring consistent with A. Unpaired t test with Welch’s correction (****p < 0.0001). **D.** Degree of lipid tail interdigitation for α = 1.00 condensate-bilayer system. **E.** Degree of lipid tail interdigitation for α = 1.00 condensate-bilayer system with tension applied in the xy-plane. Tail overlap was increased in the bilayer with tension in comparison to without tension (D). With tension, the area per lipid increases over the entire membrane. The difference between covered and protein-free regions doubled when tension was applied to the α = 1.00 system. Mann-Whitney non-parametric comparison test (****p < 0.0001). *Data was sampled every 50,000 ps and averaged per frame from two 10 µs repeat trajectories (single-sided α = 0.98) or one 20 µs trajectory (α = 1.00, α = 1.00 with tension). Symbols represent a single frame when shown. Error bars show std dev*.

## Simulation details

| $\alpha$ scaling value | total length (μs) | No. of proteins | Configuration | Tension | Related figures |
| --- | --- | --- | --- | --- | --- |
| 0.98 | 10 | 32 | double sided | n | 1, 2, 5, 6, S3, S4 |
| 0.98 | 15 | 32 | double sided | n | 1, 2, 5, 6, S1, S4 |
| 0.98 | 15 | 32 | double sided | n | 1, 2, 5, 6, S1, S4 |
| 0.98 | 10 | 36 | single sided | n | 5, S3, S4 |
| 0.98 | 10 | 36 | single sided | n | 5, S4 |
| 1.00 | 20 | 32 | double sided | n | 3, 4, S4 |
| 1.00 | 20 | 32 | double sided | y | 4, S4 |
| 1.00 | 10 | 36 | single sided | n | S3 |
| 0.9775 | 15 | 32 | double sided | n | S2 |
| 0.9775 | 15 | 32 | double sided | n | S2 |
| 0.9775 | 15 | 32 | double sided | n | S2 |

